# Conserved and species-specific STIM autoinhibition revealed by AlphaFold3-guided dissection

**DOI:** 10.64898/2026.07.27.740881

**Authors:** Panpan Liu, Xiaoqing Wu, Minhua Huang, Hongkun Wang, Pengjun Hou, Haotan He, Li Tong, Yiming Zhang, Youjun Wang

**Affiliations:** Key Laboratory of Cell Proliferation and Regulation Biology, Ministry of Education, College of Life Sciences, Beijing Normal University, Beijing, China; National Laboratory of Biomacromolecules, CAS Center for Excellence in Biomacromolecules, Institute of Biophysics, Chinese Academy of Sciences, Beijing 100101, China; Beijing Key Laboratory of Gene Resource and Molecular Development, College of Life Sciences, Beijing Normal University, Beijing, China

**Keywords:** STIM, *Caenorhabditis elegans*, FRET, calcium signaling

## Abstract

Store-operated Calcium (Ca^2+^) entry (SOCE), mediated by dynamic STIM–Orai coupling, is crucial for cellular signaling, yet the molecular architecture of STIM autoinhibition remains poorly defined. Using an AlphaFold3-guided functional validation strategy combining FRET-based biosensors, mutagenesis, and Ca^2+^ imaging, we systematically dissected the CC1–SOAR autoinhibitory interface in both human STIM1 and *C. elegans* STIM (cSTIM). Our results reveal a conserved hydrophobic core essential for maintaining the resting state in both species. Interestingly, cSTIM possesses an extended polar interaction network that strengthens autoinhibition and correlates with slower activation kinetics. Species-specific polar networks differentially modulate interface stability, providing an additional layer of regulatory tuning. This integrated approach not only validates AlphaFold3 as a powerful tool for studying dynamic regulatory interfaces but also establishes a structural framework for understanding STIM activation and disease-associated mutations, offering mechanistic insights with therapeutic implications for SOCE-related disorders.

## INTRODUCTION

Calcium ion (Ca^2+^) serves as a ubiquitous intracellular second messenger regulating various cellular processes including proliferation, differentiation, and apoptosis(Berridge *et al*, 2003; Parekh & Putney, 2005), with store-operated Ca^2+^ entry (SOCE) representing the primary route for elevating cytoplasmic Ca^2+^ levels(Chakraborty & Hasan). SOCE is orchestrated by the dynamic coupling between the ER-resident Ca^2+^ sensor STIM1 and the plasma membrane Orai1 channel protein. Dysregulation of SOCE, arising from gain- or loss-of-function mutations in STIM1, underlies diverse human pathologies including tubular aggregate myopathy, Stormorken syndrome, and immunodeficiency(Blanco *et al*; Böhm & Laporte; Lacruz & Feske, 2015; Markello *et al*, 2015; Riva *et al*), highlighting the urgent need to decipher its molecular control mechanism.

STIM protein functions as a molecular switch: its N-terminal luminal EF-SAM domain senses ER Ca^2+^ store depletion, which triggers conformational rearrangement and oligomerization that enable its translocation to ER-PM junctions and subsequent Orai1 activation(Grabmayr *et al*, 2020; Lewis; Park *et al*; Prakriya & Lewis; Qiu & Lewis, 2025; Zhang *et al*; Zhou *et al*). In the resting state, the cytosolic CC1 helix is thought to fold back toward ER and dock the SOAR/CAD domain, thereby forming an inhibitory clamp that physically occludes the Orai-binding surface(Fahrner *et al*, 2014; Ma *et al*, 2015). While this CC1–SOAR autoinhibitory interaction is supported by biochemical and mutational studies(Ma *et al*., 2015; Muik *et al*, 2011; Rathner *et al*, 2021; Shrestha *et al*, 2022), the precise residue-residue contacts, overall structural organization, and evolutionary conservation of this interface remain incompletely defined— largely because of the highly dynamic nature of full-length STIM protein and the reliance on fragment-based structural data that may not preserve native interactions(Qiu & Lewis, 2025; Rathner *et al*., 2021; Stathopulos *et al*, 2013; Stathopulos *et al*, 2008; Yang *et al*, 2012). Structural studies of the *C. elegans* STIM homolog (cSTIM) have revealed important features of its luminal EF–SAM domain and cytosolic coiled-coil region(Enomoto *et al*, 2020; Yang *et al*., 2012), yet the molecular organization of its cCC1– cSOAR interface remains largely unexplored. A cross-species comparison thus offers an opportunity to reveal essential regulatory principles by distinguishing conserved core features from species-specific adaptations. This knowledge gap hinders the development of rational therapeutic strategies targeting SOCE.

Here, we address these gaps by employing an AlphaFold3-guided functional validation strategy. We first assess the predictive capability of AlphaFold3 for the hSTIM1 CC1–SOAR interface and then systematically interrogate the functional relevance of the predicted contacts. Extending this approach to the C. elegans STIM homolog (cSTIM)(Cai, 2007; Williams *et al*, 2001), which exhibits constitutive puncta formation(Gao *et al*, 2009), suggestive of altered autoinhibition, we compare the interface architectures across species. Through this integrated approach, we identify key residues that stabilize the CC1–SOAR clamp and uncover a conserved hydrophobic interaction core that is essential for maintaining the resting state, alongside species-specific polar networks that differentially modulate autoinhibition strength and activation kinetics. These findings define the molecular architecture of the autoinhibitory interface, revealing both evolutionarily conserved determinants and lineage-specific adaptations that account for the distinct regulatory properties between hSTIM1 and cSTIM. Furthermore, our data provide mechanistic insights into how disruption of these interactions may underlie disease-associated mutations, and offer a structural framework for understanding the allosteric regulation of STIM1 activation.

## RESULTS

### AlphaFold3 modeling validates and extends the architecture of the hSTIM1 autoinhibitory interface

To resolve the molecular details of the hSTIM1 CC1–SOAR interface (domain organization depicted in Figure 1A), we used AlphaFold3 to model the Ca^2+^-bound hSTIM1 dimer. We first validated the prediction approach by modeling the previously crystallized hSOAR-MAT fragment(Yang *et al*., 2012) (hSTIM1_345– 444_ with L374M/V419A/C437T mutations; PDB: 3TEQ). The predicted structure showed high local distance difference test (pLDDT) scores (pLDDT > 70), and agreed well with the experimental crystal structure, with an overall Cα root-mean-square deviation (RMSD) of ≤ 0.8 Å (Figure S1A, left). The low RMSD values support the reliability of the predicted structures(Bagaria *et al*, 2012; Maiorov & Crippen, 1994). Consistent with previous reports that the SOAR-MAT fragment is functionally indistinguishable from wild-type SOAR(Yang *et al*., 2012), the predicted wild-type SOAR structure was nearly superimposable onto the mutant (RMSD ≤ 0.5 Å; Figure S1A, right).

**Figure 1.**
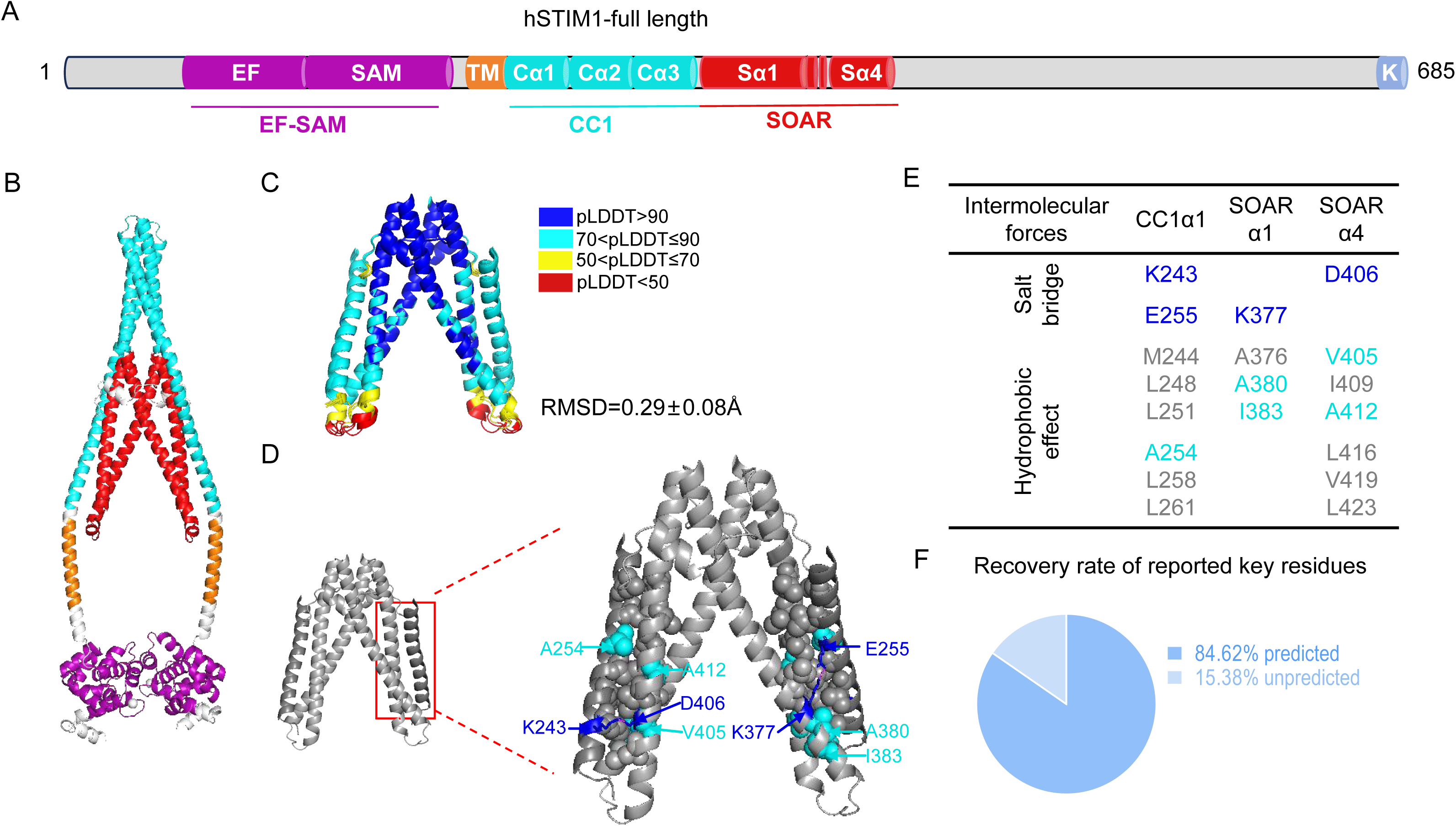
Structural insights into the Ca^2+^-bound hSTIM1 dimer and its autoinhibitory interface revealed by AlphaFold3. (A) Schematic representation of the key domains of hSTIM1: EF, EF-hand; SAM, sterile alpha motif; TM, transmembrane domain; CC1, coiled-coil 1; SOAR, the STIM-Orai activating region; K, the C-terminal Lys-rich domain. (B) Structure of Ca^2+^-bound hSTIM1 dimer predicted by AlphaFold3, with domains colored corresponding to their schematic representations in (A). (C) Predicted structure of the linked hCC1α1-hSOARL complex, with model reliability indicated by scores of predicted local distance difference test (pLDDT); the flexible linker is not shown. RMSD, the root-mean-square deviation of the five predicted configurations. (D) Close-up of the predicted hCC1 and hSOARL autoinhibitory interface, highlighting potential key interaction sites, salt bridges (blue sticks), and hydrophobic packing (gray or cyan spheres). (E) Functionally validated hydrophobic and salt-bridge interaction sites between hCC1α1 and hSOAR. Experimentally validated residues in this work: blue (See Figure S2 for complete list). (F) Accuracy of AlphaFold3 in predicting experimentally identified residues critical for maintaining the autoinhibitory state of hSTIM1.

We then modeled the Ca^2+^-bound hSTIM1_43-460_ dimer (Figures 1B and S1B). While the transmembrane (TM) and CC1 regions exhibited lower pLDDT scores—possibly reflecting inherent structural flexibility or prediction uncertainty—four of the five predicted configurations placed the TM domains in a parallel arrangement (Figure S1B), consistent with prior topological models(Jennette *et al*, 2022). Key domains displayed high conformational consistency across models: the EF-SAM monomer had an RMSD of ≤ 0.2 Å, the TM monomer showed an RMSD ≤ 0.3 Å and the SOAR dimer showed an RMSD ≤ 0.7 Å (Figure S1C), closely matching the SOAR-MAT structure (RMSD ≤ 1.4 Å) (Figure S1D). The CC1α1-SOARL complex also adopted similar conformations in all five models (Figure S1C), indicating a stable, conserved autoinhibitory clamp between the CC1α1 and SOAR. In contrast, the CC1α2-CC1α3 region displayed three distinct conformational states among the models and the pLDDT values are low, suggesting that the structure of this region may be flexible (Figure S1C).

Because the hSTIM1_238-261_ domain within CC1α1 constitutes the minimal region for SOAR docking(Ma *et al*., 2015), we next predicted the structure of the CC1α1–linker–SOARL dimer. Key regions of the complex showed high prediction confidence (pLDDT > 70), and the five replicate predictions exhibited low conformational variation (RMSD = 0.29 ± 0.08 Å; Figure 1C), supporting the reproducibility of the model.

Further analysis of the predicted interface revealed that the autoinhibitory clamp is primarily maintained by hydrophobic interactions and electrostatic contacts (Figures 1D, 1E and S2). The model recapitulates ∼85% of the functionally key residues reported to be crucial for hSTIM1 autoinhibition, including all reported hydrophobic residues (Figure 1F, and Table S1), thereby supporting the overall reliability of the predicted structure. Importantly, the model newly identifies several previously uncharacterized hydrophobic residues and two pairs of symmetric salt bridges that may contribute to the interface stability.

### Functional validation of key interactions predicted by AlphaFold3 to govern hSTIM1 autoinhibition

To investigate the functional impact of the newly predicted residues on hSTIM1 autoinhibition, we adopted a live-cell Förster resonance energy transfer (FRET) biosensor engineered to directly report conformational changes in hSTIM1 upon activation(Du *et al*, 2024; Ma *et al*., 2015) (Figure 2A, top panel). This sensor comprises two fluorescently labeled fragments of hSTIM1: hSTIM1_1-CC1_ (hSC)-ECFPΔC11 and mNGΔN5-hSOARL. In resting cells, intramolecular association maintains the two fragments in proximity, yielding a high basal FRET signal. Depletion of ER Ca^2+^ store with ionomycin (IONO), a Ca^2+^ ionophore that induces store depletion, triggered their dissociation, resulting in a rapid decrease in FRET efficiency (Figure 2A, bottom panel). This system provides a sensitive, real-time readout of hSTIM1 activation dynamics in situ.

**Figure 2.**
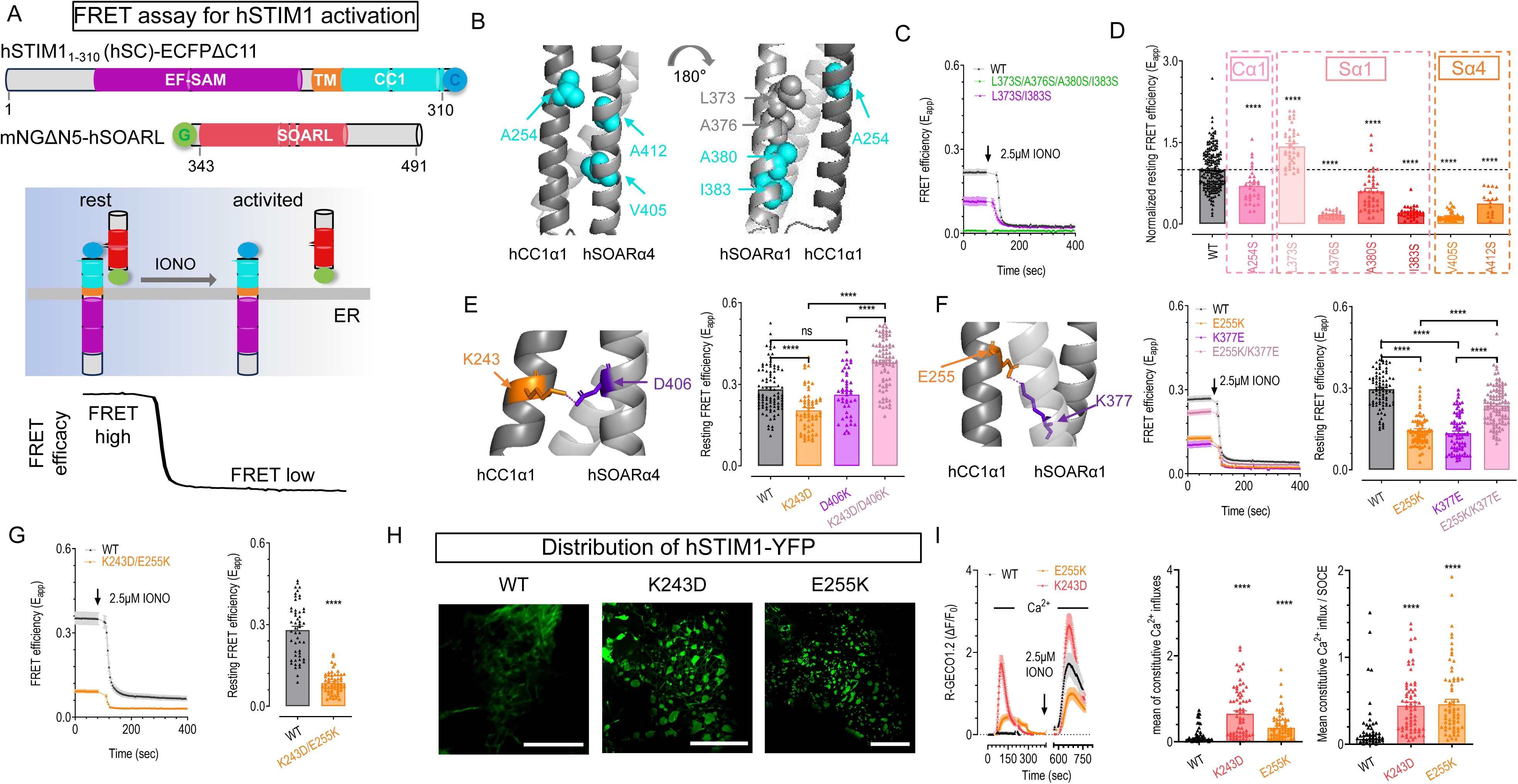
Validation of hydrophobic residues and salt bridges critical for hSTIM1 autoinhibition at the hSTIM1-CC1α1 and hSOAR interface. (A) Monitoring hSTIM1 activation by FRET. (Top) Schematics of the FRET sensor constructs: hSTIM1_1-CC1_ (hSC)-ECFPΔC11 and mNGΔN5-hSOAR1L. C, ECFPΔC11; G, mNGΔN5; (Middle) model depicting the dynamic subcellular co-redistribution of the FRET sensor pair, either at rest or upon store depletion with ionomycin (IONO) in live cells. (Bottom) Real-time FRET signal changes corresponding to the conditions above. (B-D) Validation of the predicted hydrophobic interactions via FRET. (B) Predicted hydrophobic interface between the hCC1α1 and hSOAR domains, with key residues shown as spheres. (C) Representative FRET response for WT sensor and variants carrying serine (S) substitution at the indicated hydrophobic positions. (D) Quantification of normalized resting FRET efficiency, demonstrating the effect of mutating hydrophobic residues to S. (E-F) Validating predicted salt bridges by FRET. (Left) Structure model of predicted salt bridges (stick presentation) between the hCC1α1 and hSOAR domains; (right) quantification of average resting FRET efficiency for WT and indicated charge reversal mutants. (E) The K243/D406 salt bridge (F) The E255/K377 salt bridge (G) FRET responses of WT sensor and K243D/E255K mutant. (Left), Typical traces; (right), statistics of basal FRET signals. (H) Subcellular localization of hSTIM1 variants. Representative confocal images showing the distribution of transiently expressed hSTIM1 constructs-WT, K243D, and E255K mutants in unstimulated HEK 293 cells. Scale bar, 10 µm. (I) Effect of charge reversal mutations on constitutive Ca^2+^ influx in HEK 293 cells stably expressing a red Ca^2+^ indicator, R-GECO1.2. (Left) Typical traces of R-GECO1.2 responses to extracellular Ca^2+^ addition under two conditions: at rest (reflecting constitutive Ca^2+^ entry) or after ER store depletion (reflecting SOCE). Signals corresponding to 2.5 μM IONO-induced ER store depletion phase are omitted for clarity; (Middle), mean constitutive Ca^2+^ influx; (Right) Amplitude of constitutive Ca^2+^ entry normalized to SOCE. Three independent biological replicates, at least 6 cells were examined each time. Data shown as mean ± SEM. ****, p < 0.0001, Student’s t-test.

Given the established importance of hSOARα4 in maintaining hSTIM1 autoinhibition (Figures 1E and 2B, left panel)(Horvath *et al*; Ma *et al*; Muik *et al*., 2011; Shrestha *et al*., 2022), we first examined potential novel hydrophobic interface sites in the hSOARα1 domain (Figure 2B, right panel). Comparison of FRET signals between hSC-L373S/A376S/A380S/I383S and WT hSC revealed a pronounced defect: the quadruple hSC mutant exhibited minimal basal FRET with hSOARL, with levels comparable to those observed after store depletion, indicating a near-complete loss of SOARL binding and autoinhibition at rest (Figure 2C, green trace). A double mutant, hSC-L373S/I383S, reduced the basal FRET signal by approximately half, suggesting that the other two residues A376 and A380 also substantially contribute to the hydrophobic interface (Figure 2C, purple).

We then systematically evaluated individual newly predicted hydrophobic residues by introducing single-point serine substitutions. As summarized in Figure 2D, all single-point mutants except for L373S— which lies at the periphery of the predicted hydrophobic cluster—showed a significant reduction in basal FRET, reinforcing the functional relevance of these sites in stabilizing the hCC1-hSOAR interaction.

Next, we evaluated the role of the two predicted salt bridges at the symmetric inter-subunit interface. Disruption of either salt bridge through single charge reversal mutations was expected to impair the CC1-SOAR interaction, leading to reduced FRET efficiency. As anticipated, single mutants of salt bridge pairs exhibited a decrease in basal FRET (Figures 2E and 2F, orange and purple traces). In contrast, double charge reversal mutants designed to restore complementary charges (E255K/K377E or K243D/D406K) largely recovered FRET signals, supporting the functional importance of these electrostatic interactions (Figures 2E and 2F, pink traces). Consistent with an additive effect, the hSOARL-K243D/E255K double mutant, which disrupts both salt bridges, showed a further reduction in basal FRET compared to either K243D or E255K single mutant (Figure 2G). The diminished basal FRET in these mutants reflects impaired autoinhibition and suggests constitutive activation of hSTIM1.

To assess functional consequences, we introduced the K243D or E255K mutation into full-length hSTIM1 and performed confocal and Ca^2+^ imaging. Unlike the diffuse ER-like distribution of WT hSTIM1 under resting conditions, both mutants formed constitutive puncta at the cell periphery (Figure 2H). Functionally, cells expressing hSTIM1-K243D or -E255K exhibited significant constitutive Ca^2+^ influxes, reaching approximately half the amplitude of SOCE (Figure 2I). These data collectively confirm that the two salt bridges predicted by AlphaFold3 are essential for maintaining the autoinhibitory state of hSTIM1.

Together, our results demonstrate that AlphaFold3 can reliably identify key residues that stabilize the autoinhibitory interface of hSTIM1.

### AlphaFold3 reveals a longer, more polar interface underlying slower cSTIM activation

We next applied AlphaFold3 guided FRET validation strategy to dissect the less defined activation mechanism of cSTIM (Figure 3A). Sequence alignment between hSTIM1 and cSTIM showed moderate overall identity (Figure S3), suggesting potential structural similarity. Mirroring the prediction for hSTIM1, the predicted structures of key domains of cSTIM aligned well with published ones (Figure S4A), yet a high-confidence model of the full-length protein remained elusive (Figures 3B and S4B). We therefore focused on the isolated cCC1α1-cSOAR complex. This core interface exhibited high predicted confidence, with pLDDT scores consistently above 70% and hydrophobic cluster domains exceeding 90% (Figures 3C and 3D). Structural consistency was further supported by a low RMSD (0.48 ± 0.18 Å), indicating a reliable prediction of the autoinhibitory interface.

**Figure 3.**
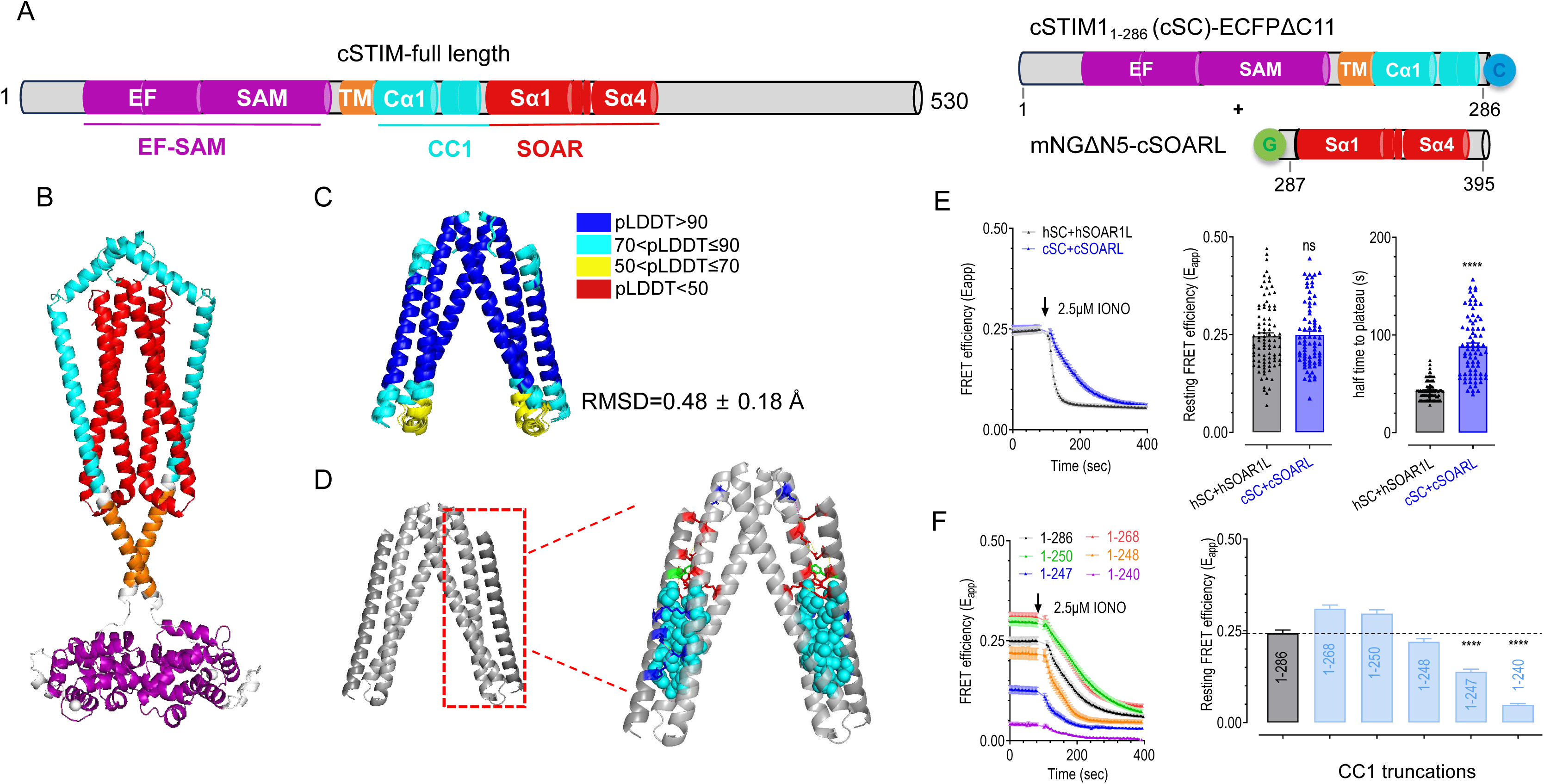
Structural insights into the Ca^2+^-bound cSTIM dimer and its autoinhibitory interface revealed by AlphaFold3. (A) Domain organization of cSTIM (left) and schematic of the FRET assay between cSTIM1_1-286_ (cSC)- ECFPΔC11 and mNGΔN5-cSOARL constructs (right). (B) Structure of Ca^2+^-bound cSTIM dimer predicted by AlphaFold3, with domains colored corresponding to their schematic representations in (A). (C) Predicted cCC1α1-cSOARL complex, with model reliability indicated by scores of predicted local distance difference test (pLDDT). RMSD, the root-mean-square deviation of the five predicted configurations. (D) Close-up view of the predicted cCC1α1-cSOARL interface, highlighting hydrogen bonds (red sticks), salt bridges (blue sticks), π-π interaction (green sticks) and hydrophobic residues (cyan spheres). (E) Activation kinetics comparison between hSTIM1 and cSTIM measured by FRET. (Left) Representative FRET responses following store depletion induced by 2.5 μM IONO; (middle) resting FRET efficiency; (right) half-time of FRET decline upon store depletion. (F) Mapping the minimal cSOAR-docking region on cCC1α1. (Left) FRET signals of truncated cSC constructs. (Right) Resting FRET efficiency quantified for each truncation variant. ****, p < 0.0001, ns, not significant, Student’s t-test. Data shown as mean ± SEM; n ≥ 10 cells from three independent experiments.

Compared to hSTIM1, cSTIM exhibited a longer predicted autoinhibitory interface, spanning residues K219 to R248 on CC1α1 (Figure S4C), which markedly exceeds the corresponding region in hSTIM1-CC1α1 (K243-L261) (Figure S2). The extended interface is characterized by additional polar interactions at its C-terminal portion, including hydrogen bonds and salt bridges (Figure 3D), implying a potentially stronger autoinhibitory constraint. Consistent with this prediction, the resting FRET between cSC and cSOAR was comparable to that of the human counterparts, but the IONO-induced FRET decay rate between cSC and cSOAR was significantly slower than that observed with the human counterparts, indicating a markedly reduced dissociation rate of cSOAR from cSC upon store depletion (Figure 3E). To define the minimal region within cCC1 (cSTIM_217-286_) required for cSOAR docking, we conducted a series of C-terminal truncation analyses. Truncation analysis further defined R248 as a critical boundary: deletions including or beyond R248 (Δ248–286) substantially reduced basal FRET, whereas truncations preserving R248 (Δ249–286, Δ251–286, or Δ269–286) did not (Figure 3F), validating that the predicted K219–R248 region defines the functional autoinhibitory interface.

### AlphaFold3 helps reveals an autoinhibitory interface with fewer hydrophobic contacts in cSTIM

We next evaluated the contribution of hydrophobic contacts to cSTIM autoinhibition (Figures 4A, 4B, and S4C). The predicted hydrophobic cluster involves residues from cSTIM-CC1α1, cSOARα1 and cSOARα4 (Figure 4C). To disrupt hydrophobic stacking, we first mutated all predicted hydrophobic residues in each of these domains (V220S/L223S/L227S/L230S/M233S in cCC1α1; A319S/I320S/V323S/L326S in cSOARα1; V350S/I354S/I357S in cSOARα4) and measured their interactions with cSOAR by FRET. All three multiple-residue mutants reduced basal FRET to levels indistinguishable from those after store depletion with IONO, indicating a near-complete disruption of cCC1-cSOAR interaction (Figure S5A), confirming that the hydrophobic core critically stabilizes the resting-state of cCC1-cSOAR complex.

**Figure 4.**
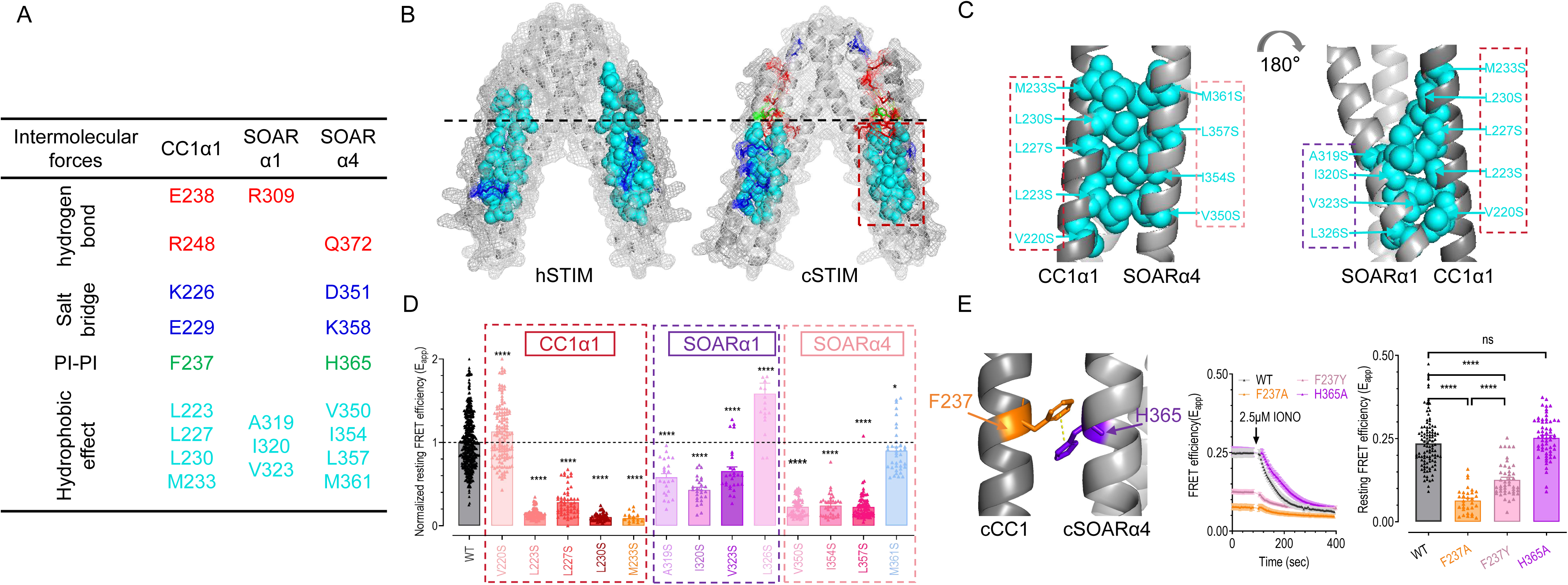
Validation of predicted cCC1-cSOAR interface crucial for cSTIM autoinhibition. (A) Summary of functionally validated cCC1α1-cSOAR interaction sites (see Figure S4C for complete list). (B) Comparison of the predicted autoinhibitory interfaces in hSTIM1 and cSTIM. Hydrophobic residues are shown as cyan spheres; polar interactions are represented as sticks (red: H-bonds; blue: salt bridges, green: π-π interaction). (C) Detailed view of the hydrophobic interface between cCC1α1 and cSOAR. (D) Quantification of normalized resting FRET efficiency for individual h-to-S point mutations. (E) Functional validation of π-π stacking pairs by FRET. (Left) Typical traces; (right), statistics of basal FRET signals. Three independent biological replicates, at least 10 cells were examined each time. *, p < 0.05; ****, p < 0.0001, ns, not significant, Student’s t-test. Data shown as mean ± SEM.

We then performed single-residue mutagenesis to assess the role of each predicted hydrophobic residue. Consistent with their peripheral location within the predicted hydrophobic cluster (Figure 4C), mutations V220S and L326S did not lower basal FRET, and M361S caused only a slight reduction (Figure 4D). In contrast, all other single-point mutants dramatically reduced basal FRET (Figure 4D), confirming that these residues are crucial for stabilizing the cCC1-cSOAR autoinhibition interface. Compared with hSTIM1, the hydrophobic cluster in cSTIM is shorter and contains fewer (11 versus 15) hydrophobic residues (Figures 1E, 4A and 4B). Overall, mutations in cSOARα1 had less severe effect than those in CC1α1 and cSOARα4, suggesting that CC1α1-cSOARα4 interaction contribute more substantially to autoinhibition. Notably, disruption of hydrophobic contacts in distinct regions each led to constitutive activation, indicating that the hydrophobic interface functions as a cooperative unit: compromising any critical node destabilizes the entire autoinhibited state.

Interestingly, a π-π interaction between F237 and H365 was also predicted. Compared with WT, the F237A mutation significantly reduced the resting FRET, whereas the milder F237Y mutation caused less pronounced impairment, indicating that this π-π stacking contributes to stabilizing the cCC1-cSOAR autoinhibition interface (Figure 4E). However, the H365A mutation showed no effect on FRET, indicating that the functional relevance of this π–π interaction may be limited. This limited impact is consistent with the location of this interaction within a polar interaction domain, where H365 is additionally predicted to engage in a polar interaction with E234 (Figure S4C). Together, these observations suggest that while the π-π stacking between F237 and H365 exists, its contribution to autoinhibition is likely context-dependent and secondary to other interactions.

### AlphaFold3-predicted additional polar interactions are critical for the enhanced cSTIM autoinhibition

After validating the hydrophobic interactions within the autoinhibitory interface, we next examined the role of predicted polar contacts. We first designed a series of charge-reversal mutations for four predicted salt bridges: K219-D351, K226-D351, E229-K358, and R248-E379 (Figures S4C, S5B-S5C and 5A-5B).

Disruption of each pair was expected to compromise autoinhibition and lower resting FRET efficiency. Consistent with the negligible functional effect seen for peripheral hydrophobic residues (hSTIM-L373, cSTIM-V220/L326; Figures 2D and 4D), single mutations of K219 and D351—predicted to form a salt bridge at the periphery of the autoinhibition interface, outside the hydrophobic cluster—also did not alter basal FRET (Figures S4C and S5B), indicating that this interaction contributes little to cSTIM autoinhibition. In contrast, the other five point mutations (K226D, E229K, K358E, R248E, and E379R) all significantly reduced basal FRET (Figures 5A, 5B and S5C, orange and purple traces).

**Figure 5.**
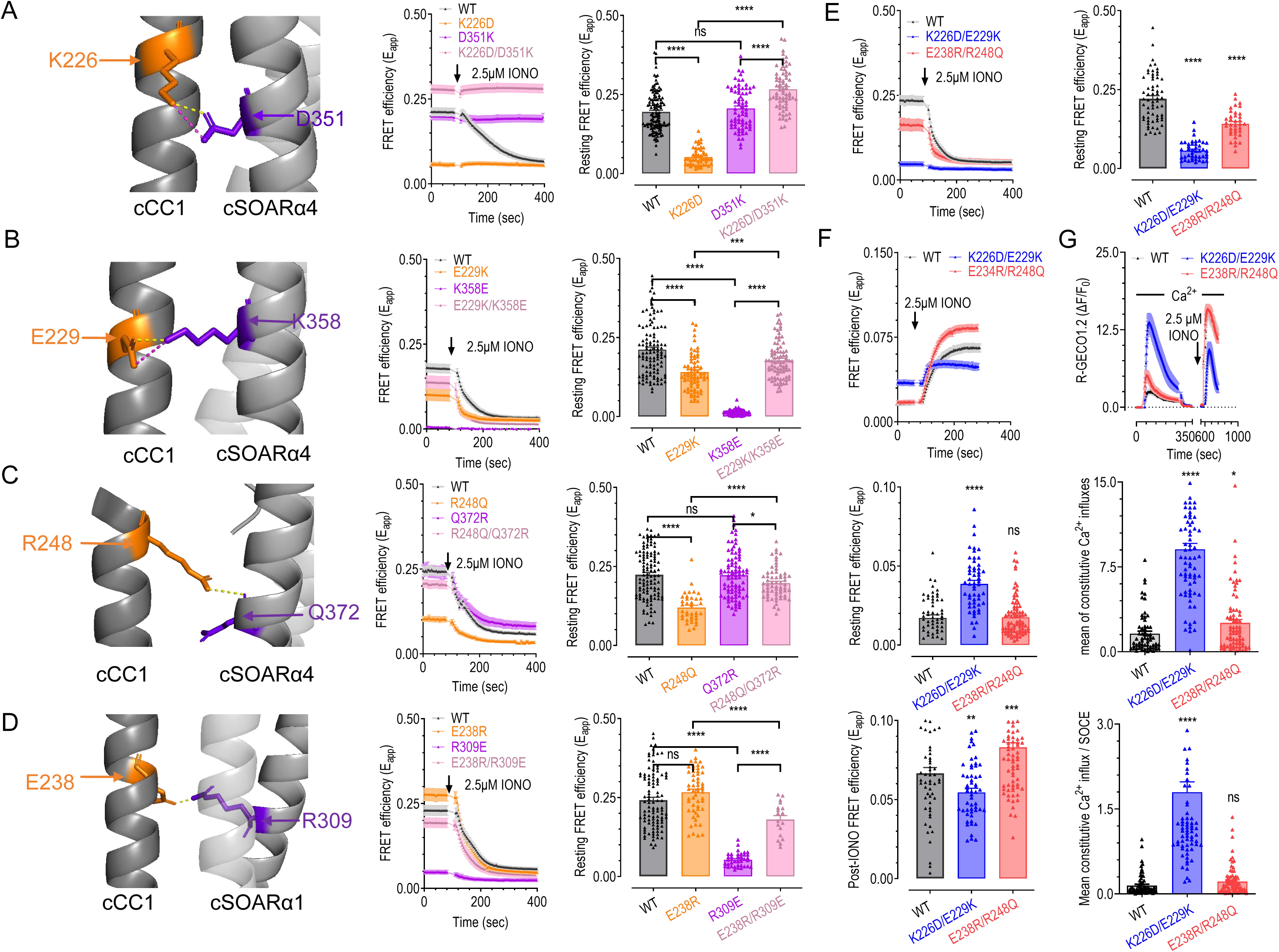
Key polar interactions act as dominant drivers of cSTIM autoinhibition. (A-D) Functional validation of individual polar pairs by FRET. For each pair, the panels show: (Left) Enlarged structural view of the specific predicted polar interaction. (Middle) Representative FRET traces. (Right) Statistics of resting FRET efficiency. Specific pairs examined. (A) K226/D351, (B) E229/K358, (C) R248/Q372, (D) E238/R309. (E) Effects of charge-reversal mutations in two predicted polar pairs on FRET responses of cCC1 variants and co-expressed cSOARL. (Left) Typical FRET traces. (Right) Statistics. (F) Impacts of polar interactions disruption on cSTIM-cOrai coupling assessed by FRET in HEK 293 cells expressing cOrai-Turquoise2. (Top) Typical traces. (Middle) Statistics of basal FRET efficiency. (Bottom) Statistics of peak FRET induced by IONO. Mutations tested, cSTIM-K226D/E229K (targeting salt bridge pairs) and cSTIM-E234R/R248Q (targeting hydrogen bond pairs). (G) Impacts of polar interactions disruption on cSTIM-cOrai mediated Ca^2+^ signals in HEK 293 cells stably expressing R-GECO1.2 with transient co-expression of cSTIM variants and cOrai. The same two mutations as those in (F) were examined. (Top) Typical traces of R-GECO1.2 responses to extracellular Ca^2+^ addition under two conditions: at rest (reflecting constitutive Ca^2+^ entry) or after ER store depletion (reflecting SOCE). Traces corresponding to 2.5 μM IONO-induced ER store depletion phase are omitted for clarity; (Middle), mean constitutive Ca^2+^ influx; (Bottom) amplitude of constitutive Ca^2+^ entry normalized to SOCE. Three independent biological replicates, at least 6 cells were examined each time. *, p < 0.05; **, p < 0.01; ****, p < 0.0001, ns, not significant, Student’s t-test. Data shown as mean ± SEM.

The corresponding double charge-reversal mutants for two of the salt bridge pairs (K226-D351 and E229-K358) largely restored FRET signals to near WT levels (Figures 5A and 5B, pink traces), confirming that these electrostatic interactions are critical for stabilizing the autoinhibitory state. Notably, the reciprocal charge-swap double mutant R248E/E379R, unlike the other rescued pairs, failed to restore FRET (Figure S5C), suggesting that the R248-E379 interaction may depend on additional structural constraints, such as the auxiliary hydrogen bond between R248 and Q372 (Figure 5C).

We further examined the five predicted polar interactions (Figures 5C, 5D, S5D–S5F) by assessing the effects of reciprocal point mutations on basal FRET. Mutations E234R, Q241A and Q372A had negligible effects, indicating that the E234-R309 and Q241-Q372 polar pairs contribute little to cSTIM autoinhibition. Similarly, the E234H single mutant showed no change in FRET, and the double mutant E234H/H365E failed to rescue impairment caused by H365E, confirming that E234-H365 polar interaction also plays a minor role. The reduction of basal FRET by H365E is likely caused by disruption of π-π stacking between H365 and F237 (Figure 4E). In contrast, while the single mutants E238R and Q372R had no effect on basal FRET, the corresponding double reciprocal mutant (E238R/R309E and R248Q/Q372R) largely restored impairment caused by R309E and R248Q, respectively, demonstrating that the E238-R309 and R248-Q372 polar pairs are important for maintaining cSTIM autoinhibition. To further assess the relative contribution of different polar interaction types, we compared the effects of disrupting the two validated salt-bridges versus hydrogen-bonds using double mutants in cSTIM-CC1α1. Breaking both salt bridges (K226D/E229K) caused a greater reduction in basal FRET than disrupting the hydrogen bond pairs (E238R/R248Q) (Figure 5E), suggesting that electrostatic interactions play a more dominant role in maintaining the cSTIM autoinhibition.

To evaluate the effects of distinct polar interactions in cSTIM-cOrai coupling, we performed FRET and Ca^2+^ imaging assays on full length cSTIM constructs carrying K226D/E229K and E238R/R248Q double mutations. In control experiments, HEK 293 cells co-expressing WT cSTIM and cOrai showed low basal FRET that increased upon IONO stimulation, reflecting store-depletion-induced coupling. In contrast, the cSTIM-K226D/E229K mutant exhibited elevated basal FRET with only a slight increase upon IONO treatment, indicative of constitutive cSTIM activation and pre-coupled association with cOrai. The cSTIM -E238R/R248Q mutants showed negligible effects, indicating that hydrogen bonds contribute minimally to cSTIM-cOrai pre-coupling (Figure 5F). We next quantified constitutive Ca^2+^ entry using Ca^2+^ imaging. When normalized to SOCE, the amplitude of constitutive Ca^2+^ influx corresponded to 100% activation for cSTIM-K226D/E229K and 22% for cSTIM-E238R/R248Q, further demonstrating that the effect of electrostatic interactions exerts a significantly greater effect than hydrogen-bonds on cSTIM activation (Figure 5G). Notably, the E238R/R248Q mutants displayed IONO-induced peak FRET and SOCE amplitudes slightly higher than WT (Figures 5F and 5G), though the underlying mechanism remains to be investigated.

## DISCUSSION

Using an AlphaFold3-guided functional validation strategy, we systematically identified key residues stabilizing the STIM autoinhibitory interface, revealing a conserved hydrophobic core alongside species-specific polar networks that differentially modulate autoinhibition strength and activation kinetics. Specifically, the hydrophobic core is conserved between hSTIM1 and cSTIM and is required for maintaining the autoinhibited state in both proteins. In contrast, polar interactions showed substantially lower conservation and contributed differently to interface stability in the two species. Notably, cSTIM contains an expanded network of polar interactions that strengthens the CC1–SOAR association and correlates with slower activation following store depletion. These observations indicate that while the overall architecture of the STIM autoinhibitory interface is conserved, species-specific variations in polar interactions provide an additional layer of regulatory tuning (Figure S6).

More broadly, our study highlights the utility of AlphaFold3 for investigating regulatory protein interfaces that are challenging to resolve experimentally. Although AlphaFold3 has demonstrated remarkable success in predicting protein structures and protein complexes(Liu *et al*, 2025), its application to dynamic intramolecular regulatory interactions remains less explored. Here, AlphaFold3 not only recapitulated previously established features of the hSTIM1 inhibitory clamp but also identified numerous functionally important interactions that were subsequently validated by mutagenesis, FRET measurements, and Ca^2+^ imaging. Together with recent studies demonstrating the ability of AlphaFold3-based approaches to identify protein–protein interaction interfaces(Bryant *et al*, 2022; Lee *et al*, 2024), our findings suggest that structure prediction can serve as an effective hypothesis-generating platform for dissecting both intramolecular and intermolecular regulatory mechanisms. In the context of SOCE, it will be particularly interesting to determine whether similar approaches can help resolve the molecular interface between activated STIM and Orai, which remains elusive despite extensive investigation.

### Limitations of the study

Several limitations should be acknowledged. First, our analyses focused primarily on the CC1α1–SOAR interface and do not provide a complete description of all structural determinants governing STIM autoinhibition. Previous studies have implicated additional regions, including the CC1α2–CC1α3 domains(Qiu & Lewis, 2025), SOAR dimerization interfaces(Yang *et al*., 2012; Zhou *et al*, 2015), and the SOAR apex region(Höglinger *et al*, 2021; Qiu & Lewis, 2025), in regulating STIM activation and Orai coupling. Owing to the relatively low confidence of AlphaFold3 predictions for these regions, they were not examined further here. Future studies integrating complementary structural and functional approaches will be required to determine how these elements cooperate with the CC1α1–SOAR interface to regulate the transition between resting and activated states. Second, although extensive mutational analyses support the functional importance of the predicted interface residues, our data do not directly demonstrate the physical interactions proposed in the structural models. High-resolution structures of the autoinhibitory CC1-SOAR complex remain unavailable, largely due to the dynamic nature of this interaction. Nevertheless, the experimentally validated CC1–SOAR models presented here offer a starting point for future structural studies — whether by crystallography, cryo-EM, integrative structural approaches, or orthogonal techniques capable of probing residue-level proximity in situ, such as site-specific crosslinking and photo crosslinking strategies.

## RESOURCE AVAILABILITY

### Lead contact

Requests for further information and resources should be directed to and will be fulfilled by the lead contact, Youjun Wang.

### Materials availability

This study did not generate new unique reagents. All plasmids generated in this study are available from the lead contact with a completed materials transfer agreement.

### Data and code availability

- All data reported in this paper will be shared by the lead contact upon request.
- This paper does not report original code.
- Any additional information required to reanalyze the data reported in this paper is available from the lead contact upon request.

## ACKNOWLEDGMENTS

This work was supported by the National Natural Science Foundation of China (W2411015), the Ministry of Science and Technology of China (2023YFA1801902), and the Fundamental Research Funds for the Central Universities. We acknowledge the Microscopy Core Facility of College of Life Sciences, Beijing Normal University, for assistance with microscopy.

## AUTHOR CONTRIBUTIONS

Y.W. and Y.Z. conceived the project, supervised the study, and coordinated the overall research efforts. P.L. carried out the majority of the experimental work, including plasmid construction, live-cell Ca²⁺ imaging, FRET assays, and super-resolution imaging, and was responsible for data acquisition, organization, and statistical analysis, with input from other authors. X.W., M.H., and H.W. contributed to plasmid construction, while H.H. and P.H. assisted with part of this work. L.T. provided technical support for imaging equipment. P.L. drafted and revised the manuscript, and Y.W. revised and finalized it. All authors have read and approved the final version of the manuscript.

## STAR★METHODS

### KEY RESOURCES TABLE

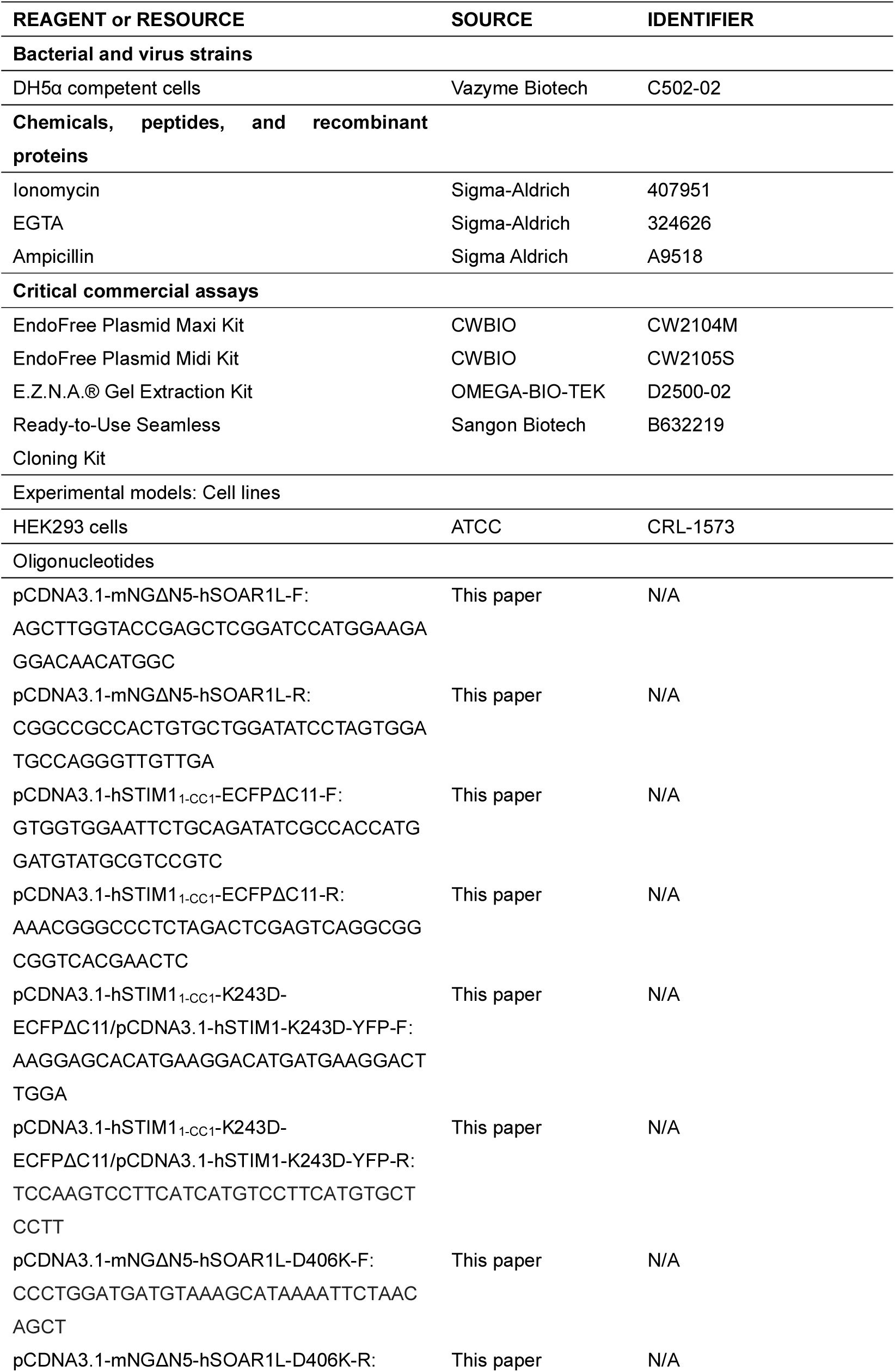
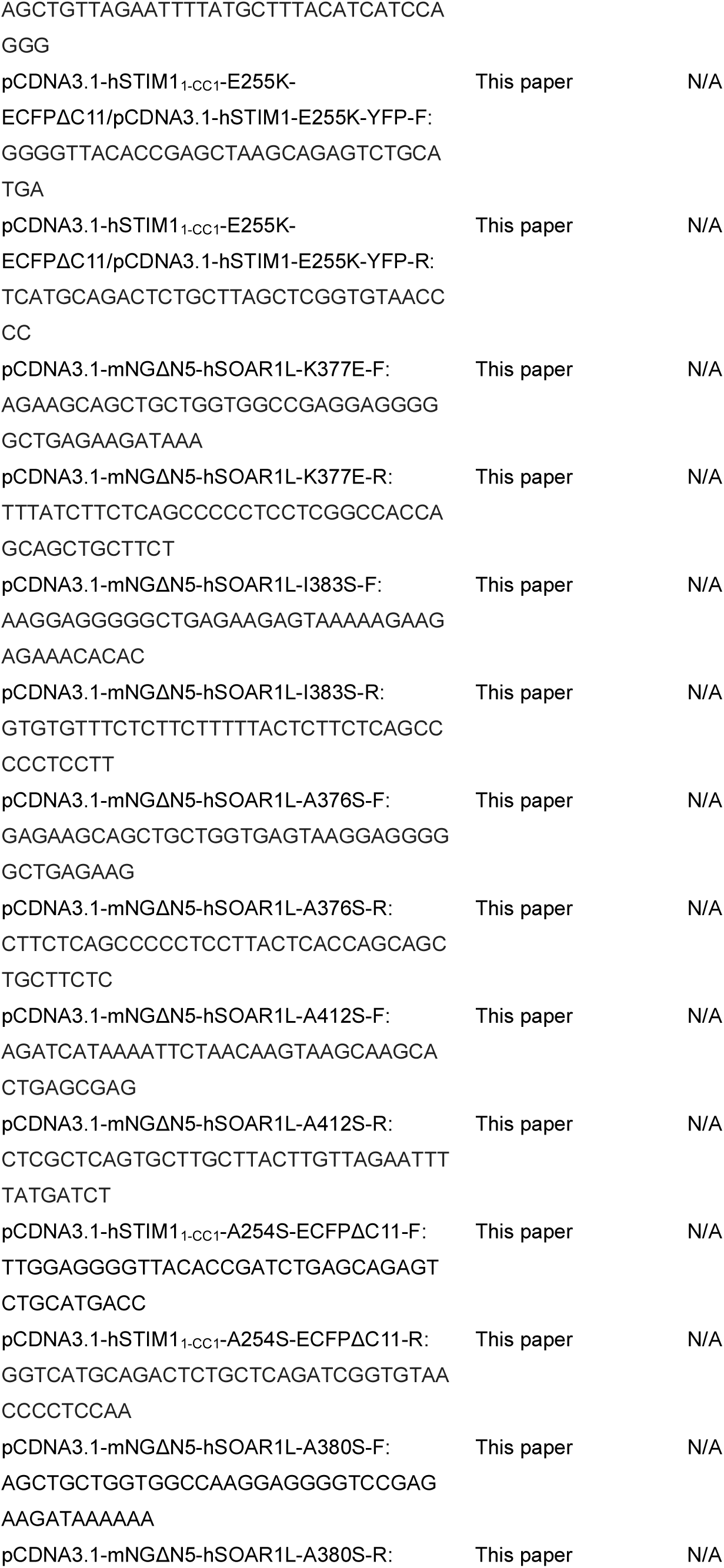
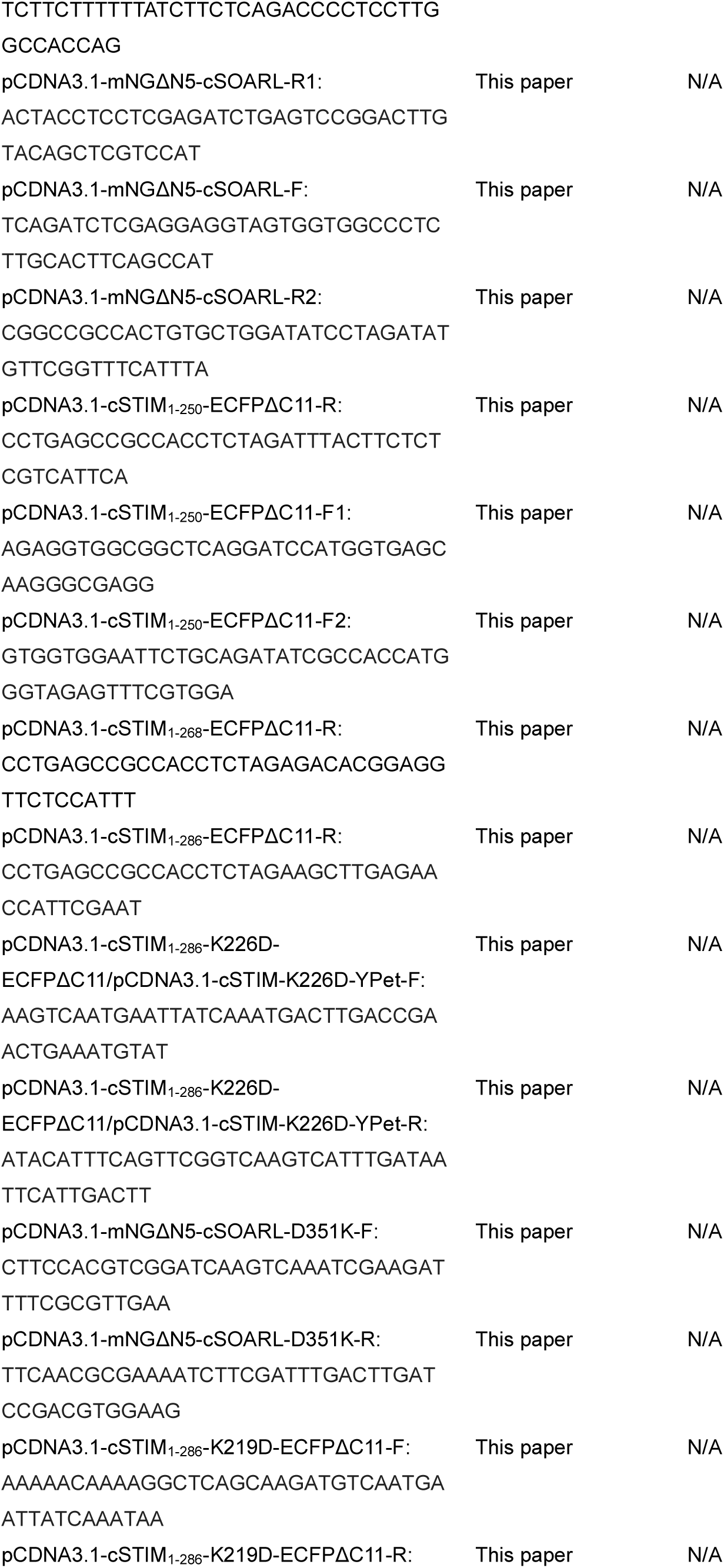
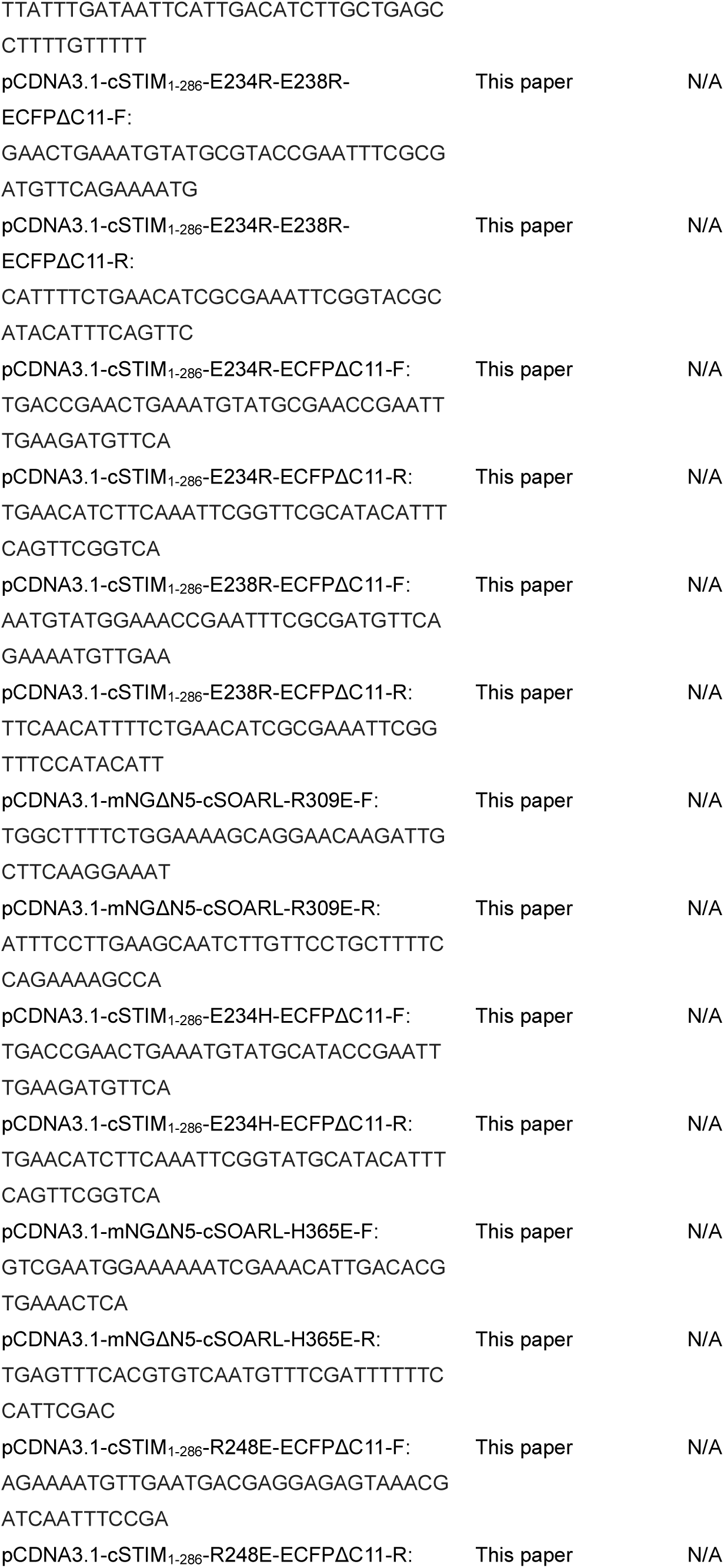
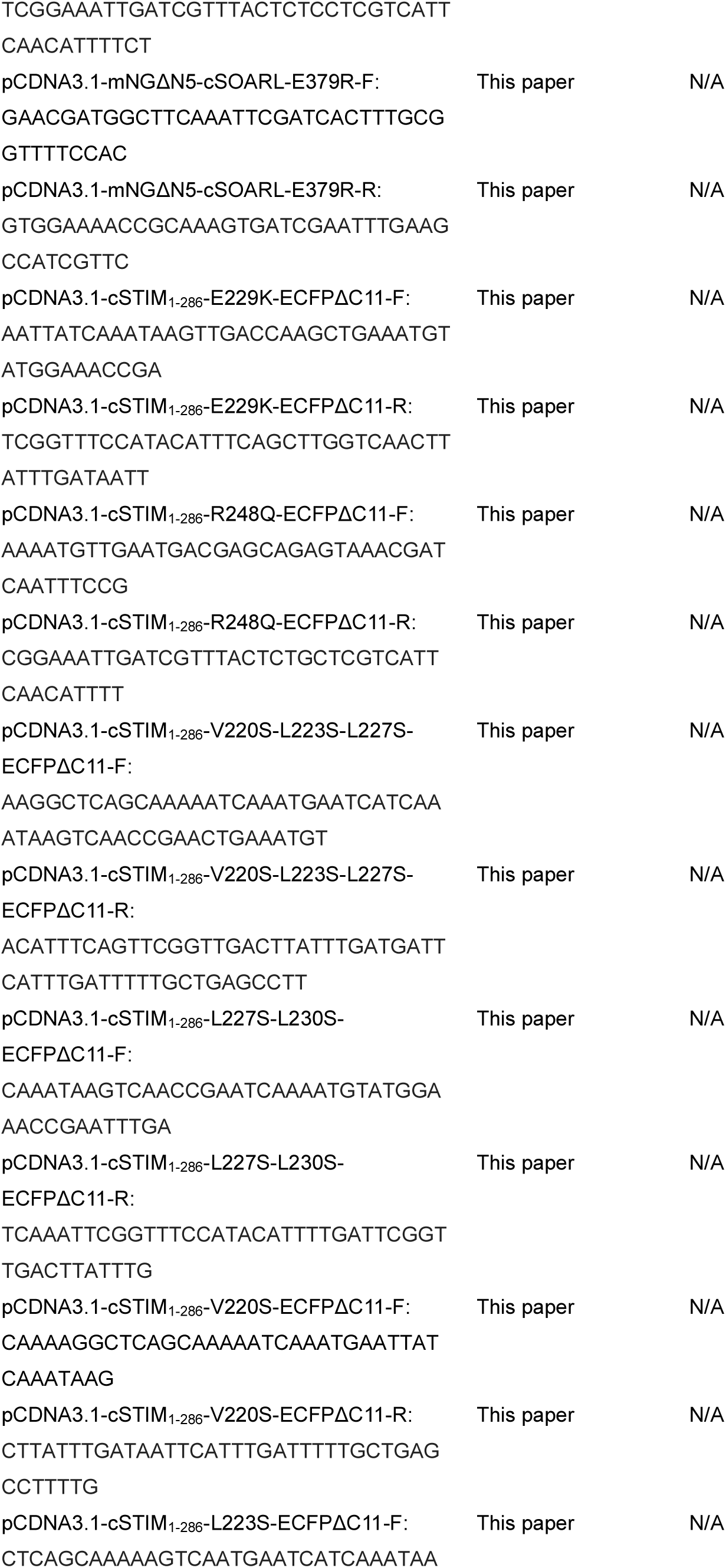
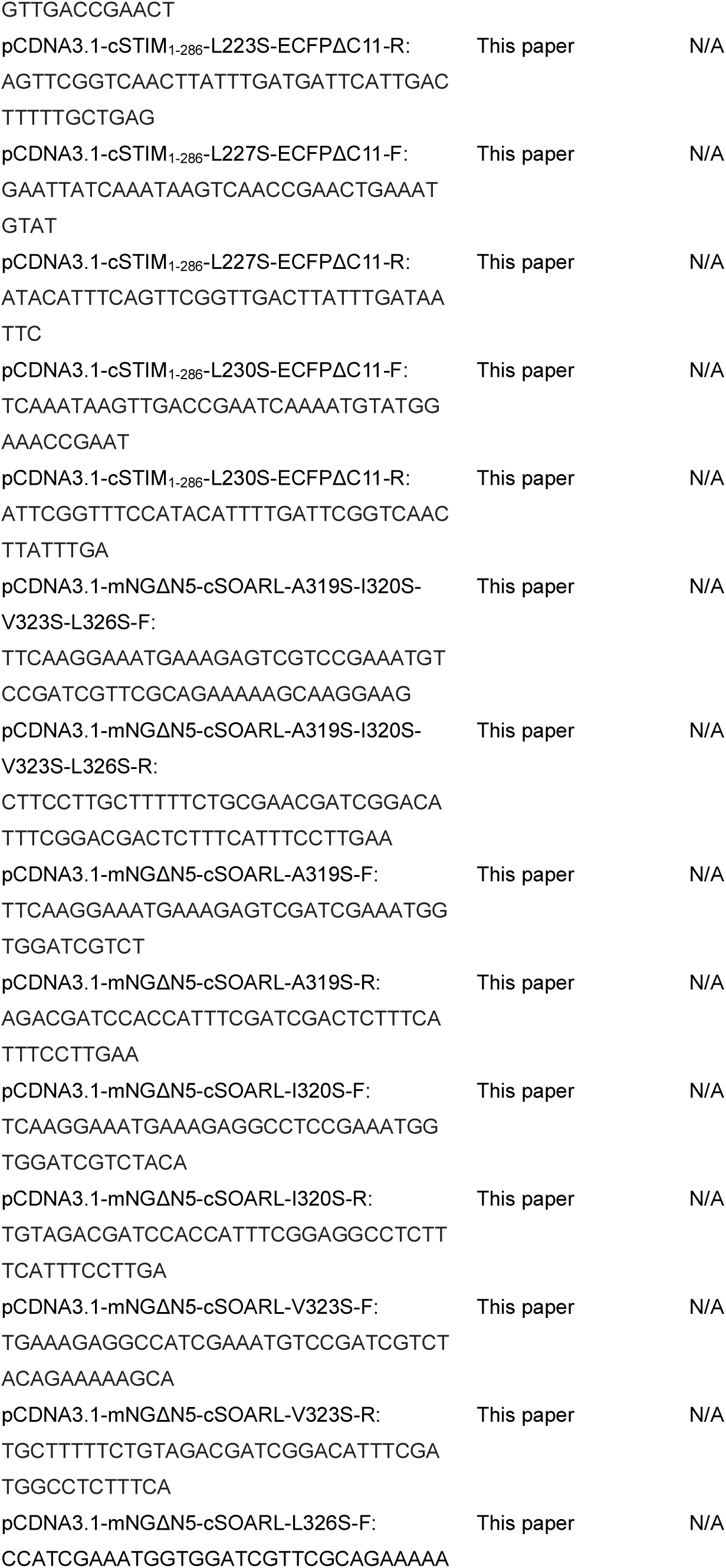
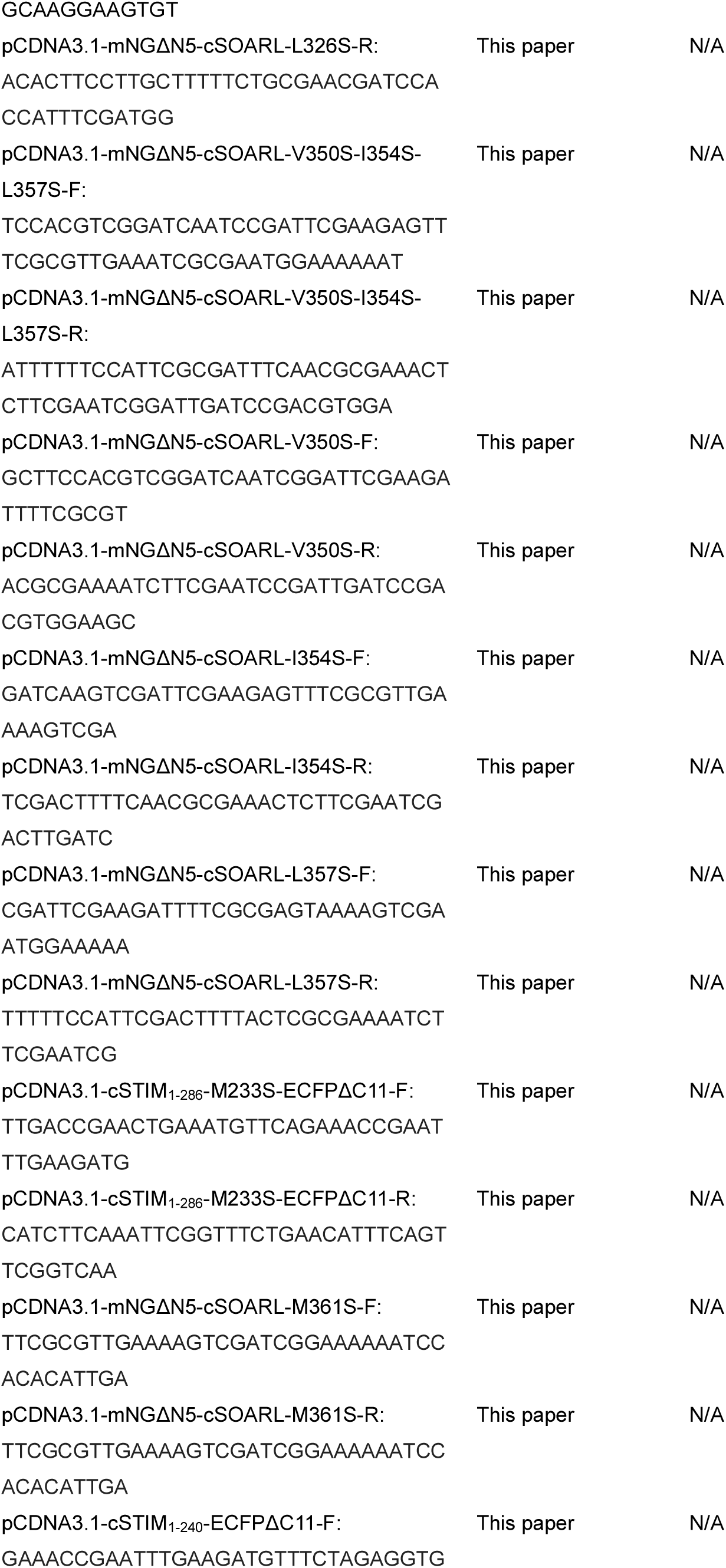
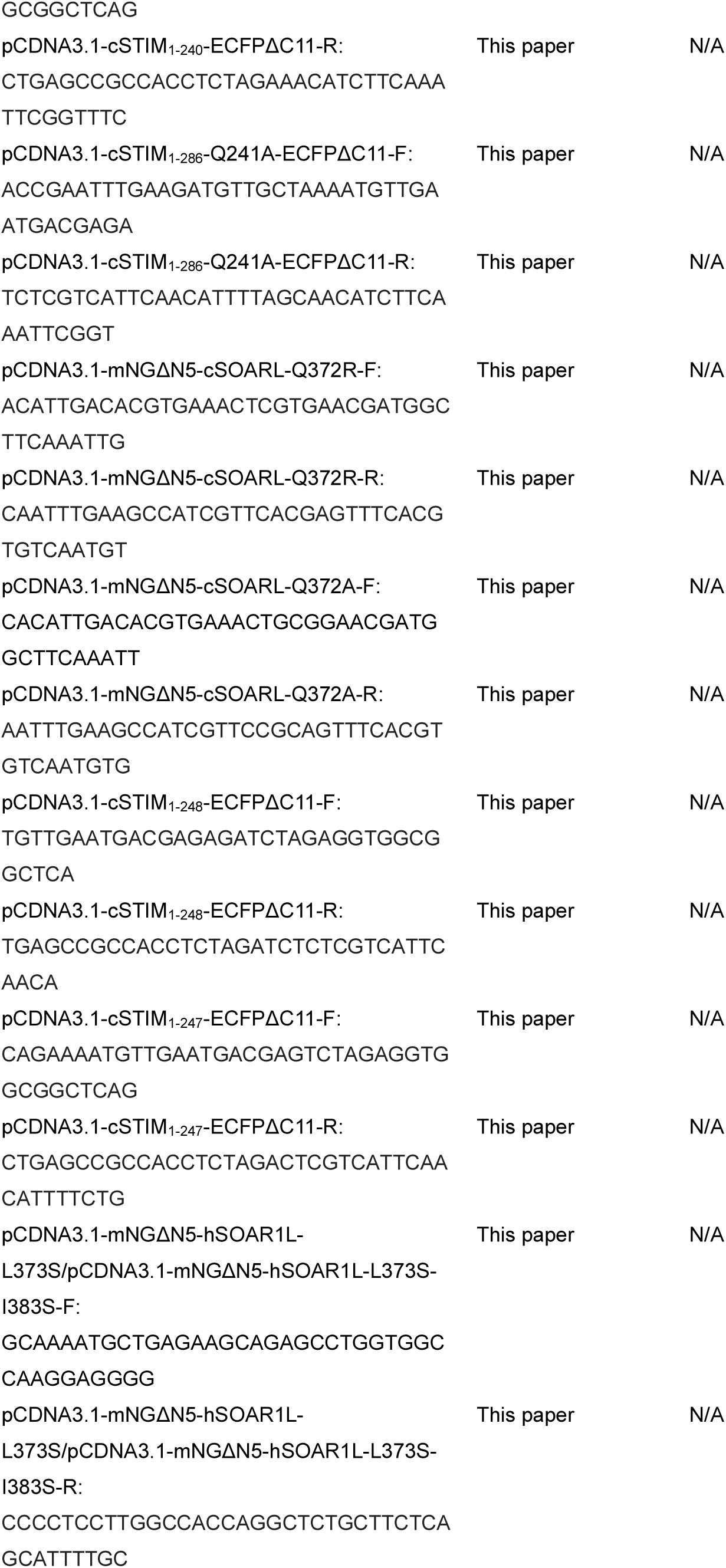
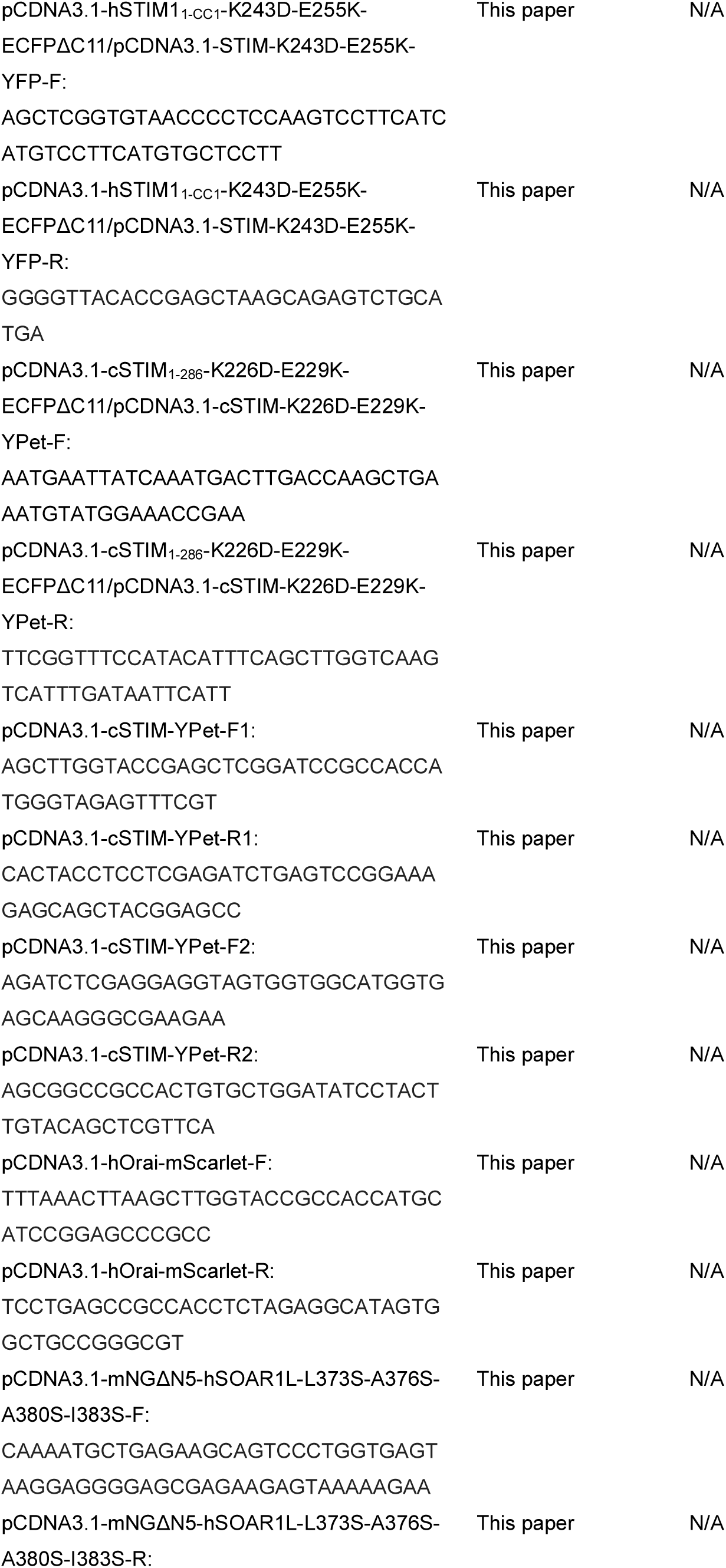
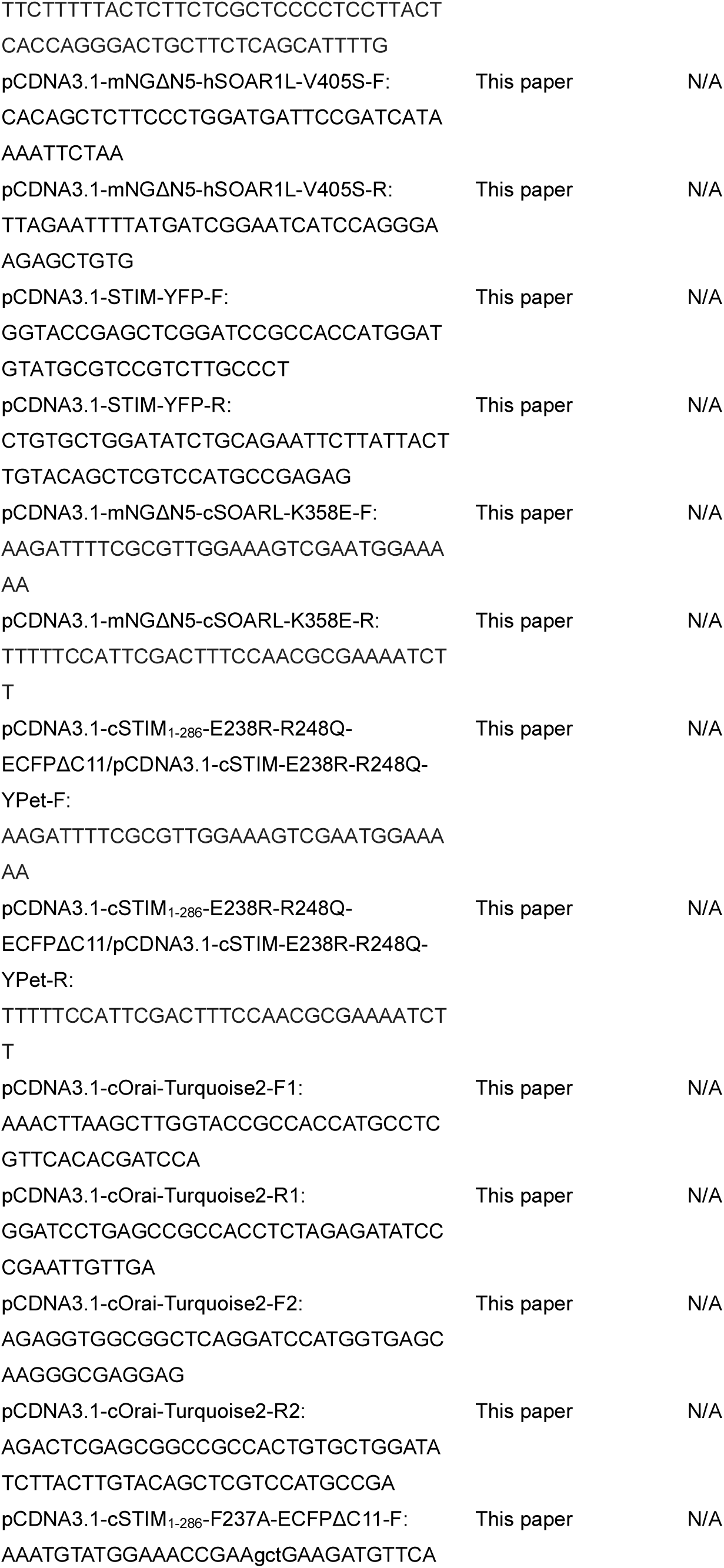
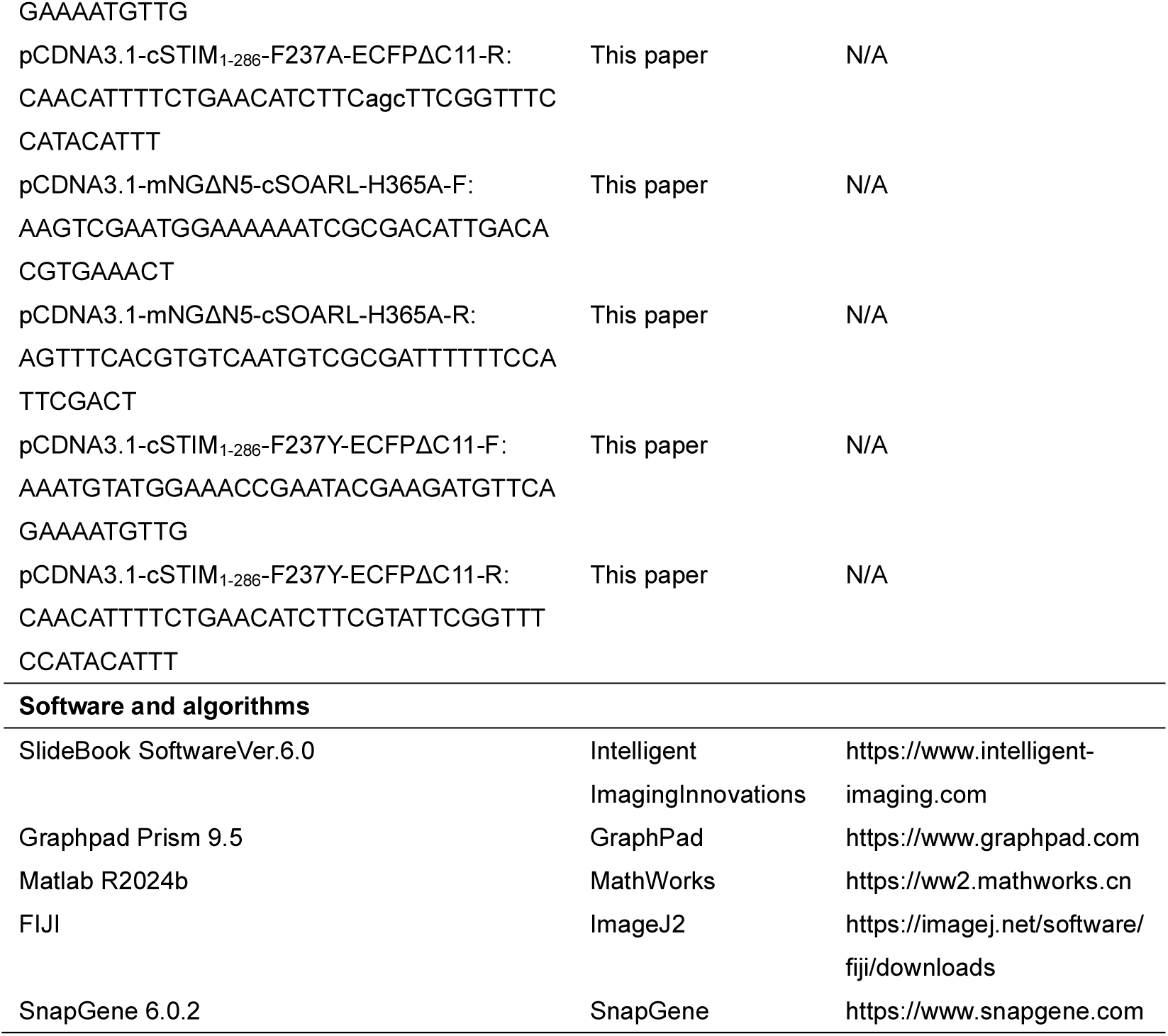

### EXPERIMENTAL MODEL AND STUDY PARTICIPANT DETAILS

#### Cell line

HEK 293 cells (ATCC, Cat. # CRL-1573) were cultured in Dulbecco’s modified Eagle’s medium (DMEM; Cytiva) with 1x penicillin–streptomycin and 10% serum standard. All cells were cultured at 37°C with 5% CO_2_.

### METHOD DETAILS

#### Cell culture and transfection

HEK 293 cells (ATCC, Cat. # CRL-1573) were maintained in Dulbecco’s modified Eagle’s medium (DMEM; Cytiva, Cat. # SH30243.01, Lot. # AL30836864) supplemented with 10% serum standard (Xrbiotech, REF. # XRSS10-500, Lot. # 001HM2444) and 1% penicillin–streptomycin (Gibco, REF. # 15140-122), at 37°C in a 5% CO_2_ atmosphere. All transfections were carried out by electroporation using the Gene Pulser Xcell system (Bio-Rad, Hercules, CA, USA). Cells were electroporated in 0.4 mL of Opti-MEM medium (Gibco, REF. # 31985-070, Lot. # CC16242) in 4 mm cuvettes with a square wave pulse (180 V, 25 ms) and then seeded onto round coverslips. After transfection, cells were cultured in serum-free Opti-MEM for 45 minutes, followed by incubation in the complete DMEM for 16-18 hours.

#### Plasmids Construction

To generate human hSTIM1 mutants, the corresponding DNA sequences were amplified via PCR using plasmids encoding hSTIM1_1-CC1_-ECFPΔC11 or mNeonGreenΔN5 (mNGΔN5)-hSOARL as templates(Du *et al*). The coding sequence (CDS) of cSTIM (NCBI: NP_741074.1) and cOrai (NCBI: NP_ 001254834.1) were synthesized and incorporated into a pCDNA3.1(+) vector by Ruibiotech (Beijing, China). Fragments and mutants of cSTIM or hSTIM1 were subsequently amplified from construct. After PCR, the fragments were reassembled and inserted into the a linearized pCDNA3.1 vector backbone-digested with appropriate restriction enzymes-using a Ready-to-Use Seamless Cloning Kit (B632219, Sangon Biotech, Shanghai, China). All primers used for generating the corresponding mutations were included in Table S2. All constructs and mutations were verified by DNA sequencing.

#### Structure prediction using AlphaFold3

Comparative structural modeling of full-length hSTIM1 (UniProt: Q13586), cSTIM (UniProt: G5EF60) were performed using AlphaFold3. For each protein, an ensemble of five models was generated. These models were ranked based on comprehensive confidence metrics, including per-residue pLDDT scores, and domain-specific reliability estimates. Additionally, to dissect domain-specific interactions and folding, we performed separate structural predictions for key isolated regions (e.g., the EF-SAM luminal domain, the CC1-SOAR cytosolic segments) of both proteins. All final models were exported for visualization and comparative analysis using PyMOL (The PyMOL Molecular Graphics System, Version 2.0 Schrödinger, LLC). This integrated in silico approach provided a high-confidence structural framework for both orthologs, enabling a direct comparison of their domain architecture, inter-domain interfaces, and regions of conformational flexibility. The resulting models form the basis for hypothesizing conserved and divergent features in their autoinhibitory mechanisms.

#### Analysis of internal interaction sites in STIM using Pymol

Structural visualization and interaction analysis of the predicted STIM model were performed using the molecular graphics software PyMOL (version 3.1.6.1, Schrödinger, LLC). First, the selected AlphaFold3 predicted model was imported into the software. To clearly display domain organization, the “cartoon” mode was used to visualize the protein backbone, with distinct coloring applied to differentiate key functional domains such as EF-SAM, CC1, and SOAR/CAD.

The study systematically identified four types of non-covalent interactions: hydrogen bonds, salt bridges and π-π interaction: detected using PyMOL’s built-in find command, with manual validation of bond lengths. Hydrophobic Interactions: Potential interaction interface residues were preliminarily screened by running a Python script (Interface.py), which identifies atoms with inter-domain or inter-chain distances of less than 5 Å. Represented by defining typical hydrophobic residues (Ala, Gly, Val, Ile, Leu, Phe, Met, Pro, Trp) and further filtering for side-chain atom clusters located at interfaces while excluding backbone atoms (Cα, N, O). All identified key interaction residues were highlighted in “stick” (polar interaction) or “sphere” (hydrophobic interaction) mode and superimposed onto the semi-transparent overall protein structure to generate a comprehensive interaction map.

#### Förster resonance energy transfer (FRET) imaging

In this study, FRET was measured using the fluorescent protein pair ECFPΔC11 (excitation 438 ± 12 nm, emission 483 ± 12 nm) and mNGΔN5 (excitation 500 ± 12 nm, emission 542 ± 13.5 nm), as well as the Turquoise2 (excitation 434 ± 12 nm, emission 474 ± 15 nm) and YPet pair (excitation 517 nm, emission 530 nm). Take ECFPΔC11 and mNGΔN5 FRET pairs as an example, during calibration, the corrected FRET signal (FRETc) was calculated from bleed-through–corrected raw signals using the following equation: FRETc=F_raw_-F_d_/D_d_×F_ECFPΔC11_-Fa/Da×F_mNGΔN5_, where F_d_/D_d_ denotes the experimentally determined bleed-through coefficient of ECFPΔC11 into the FRET channel (0.85), and F_a_/D_a_ represents the bleed-through coefficient of mNGΔN5 into the FRET channel (2.8). To account for variations in protein expression, the FRET signal was normalized to the donor fluorescence, yielding N-FRET (normalized FRET): N-FRET= FRETc/ F_ECFPΔC11_. The apparent FRET efficiency (Eapp) between the two proteins was then calculated as: Eapp=N-FRET/(N-FRET+G), with G being a system-specific correction factor (set to 7.5 in this study). It is obtained with partial mNGΔN5 photo-bleaching method: G = (FRETc-FRETc^post^)/(F_ECFPΔC11_^post^-F_ECFPΔC11_), where FRETc^post^ and F ^post^ correspond to FRETc and F_ECFPΔC11_ values after partial photo-bleach of mNGΔN5.

For functional imaging, mNGΔN5-hSOARL and hSC-ECFPΔC11 (or mNGΔN5-cSOARL and cSC-ECFPΔC11) were transiently co-expressed in HEK293 cells. Experiments were performed in imaging buffer (107 mM NaCl, 7.2 mM KCl, 1.2 mM MgCl_2_, 1 mM CaCl_2_, 11.5 mM glucose, 20 mM HEPES-NaOH and 0.1% BSA) containing 2 mM Ca^2+^ (pH 7.2). Fluorescence signals from the ECFPΔC11, mNGΔN5, and FRET channels were recorded at 2s intervals. After 30 time points, the bath solution was exchanged for a Ca^2+^-free solution supplemented with 2.5 µM IONO to deplete endoplasmic reticulum Ca^2+^ stores, thereby allowing observation of FRET changes before and after store depletion. All recordings were carried out at room temperature.

Fluorescence data from regions of interest were exported from SlideBook 6.0.23 software and processed using Matlab 2024b to calculate system-independent apparent FRET efficiency. Representative curves from at least three independent experiments are shown, with data presented as mean ± standard error of the mean (SEM). Figures were generated using Prism 9.5.1 software.

#### Ca^2+^ imaging in living cells

All Ca^2+^ imaging experiments were conducted by following our previous procedures(Gu *et al*, 2025). In brief, Cells seeded on the round coverslips were incubated in imaging buffer. Time-lapse images were performed at room temperature using a ZEISS observer Z1 imaging system controlled by SlideBook v.6.0.23 (Intelligent Imaging Innovations, Inc.). Ca^2+^ signals from HEK293 cells cotransfected with a red Ca^2+^ indicator, R-GECO1.2(Pathak *et al*), were indicated every 2 s using a TxRed-A-Basic-000 filter set, and the fluorescent intensity changes (ΔF/F0) of the corresponding regions was exported for analysis using Matlab 2024b. Representative traces from at least three independent experiments are shown as mean ± SEM.

#### Confocal microscopy

Images were undertaken with a ZEISS LSM880 system equipped with 63x oil objective (NA 1.4), controlled via ZEN 2.1 software. mNGΔN5 was excited by 488 nm laser, and the emission collected at 470–540 nm. mScarlet were excited by 543 nm laser, and the emission collected at 590–690 nm. The thickness of the optical slice is 1 µm. The resulting images taken at the footprints of cells were exported, and analyzed using Image J software. All experiments were repeated at least three independent experiments, and the representative data were shown.

## QUANTIFICATION AND STATISTICAL ANALYSIS

All quantitative data are presented as mean ± SEM from at least three independent biological replicates. Statistical significance was assessed using unpaired Student’s t-tests and paired Studen’s t-tests implemented in Matlab R2024b. *P* value < 0.05 was considered statistically significant.

